# Inferring the composition of the blood plasma proteome by a human proteome distribution atlas

**DOI:** 10.1101/2024.05.07.592979

**Authors:** Erik Malmström, Simon Hauri, Tirthankar Mohanty, Aaron Scott, Christofer Karlsson, Carlos Gueto-Tettay, Emma Åhrman, Shahab Nozohoor, Bobby Tingstedt, Sara Regner, Peter Elfving, Leif Bjermer, Andreas Forsvall, Alexander Doyle, Mattias Magnusson, Ingrid Hedenfalk, Päivi Kannisto, Christian Brandt, Emma Nilsson, Lars B Dahlin, Johan Malm, Adam Linder, Lars Malmström, Emma Nimeus, Johan Malmström

## Abstract

The plasma proteome is maintained by the influx and efflux of proteins from surrounding organs and cells. To quantify the extent different organs and cells contribute to the plasma proteome composition, we developed a mass spectrometry-based proteomics strategy to infer the origin of proteins detected in human plasma in health and disease. First, we constructed an extensive human proteome atlas from 18 vascularized organs and the most abundant cell types in blood. Second, the atlas was interfaced with previous RNA/protein atlases to objectively define proteome wide protein-organ associations to enable both the inference of origin and the reproducible quantification of organ-specific proteins in plasma. We demonstrate that the resource can determine disease-specific quantitative changes of organ-enriched protein panels in three separate patient cohorts with infection, pancreatitis, and myocardial injury. The strategy can be extended to other diseases to advance our understanding of the processes contributing to plasma proteome dynamics.

## Introduction

Systems biology type analysis of tissue and cell proteomes can reveal protein abundance and post-translational modification landscapes in tissues and cells. Such information is critical to increase the understanding of how tissue proteomes are maintained and how the tissue-specific combination of proteins enables specialized organ functions and regenerative capabilities. The ongoing cataloguing of the human proteome has revealed that tissues produce specific protein repertoires consisting of both tissue-enriched proteins and proteins that are shared across many different tissues. These protein repertoires orchestrate the tissue-specific and basic cellular functions performed in different organs and organ systems [1–3]. In contrast to tissues and cells, there is no protein synthesis in the blood plasma. Rather the composition of blood plasma proteome is maintained by the influx and efflux of proteins from surrounding cells and organs.

Based on origin and function, previous reports have shown that proteins the plasma proteome can broadly be categorized into two predominant classes ([4].). One class contains abundant proteins produced by the liver and is involved in the principal functions of plasma, such as providing colloid osmotic pressure and maintaining hemostasis through the complement and coagulation systems. Examples are human serum albumin, apolipoproteins, acute phase proteins and proteins of the complement and coagulation system. The other class, more numerous but typically less abundant, are tissue proteins that do not contribute to the principal functions of blood plasma ([4]). Diseases or injuries can induce damage to cells or tissues leading to altered abundance levels of tissue proteins in body fluids, such as plasma, that can be used as protein biomarkers for disease diagnosis and prognosis. Prominent examples are cardiac troponins (cTN) and pancreatic alpha-amylase where increased plasma concentrations are used to diagnose acute myocardial infarction (AMI) [5] and pancreatitis [6] respectively. Previous work has demonstrated that the plasma levels of tissue-enriched protein panels change in animal models are impacted during disease and can be used to assess organ dysfunction ([7]).

However, systematic investigations of the properties and dynamics regarding the infiltration of tissue/cellular proteins in the plasma during health and disease at a proteome-wide scale in human subjects is currently missing.

Here we used data-independent acquisition (DIA) mass spectrometry to create a proteome distribution atlas in healthy humans covering the major human tissues and blood cells. Further, we added data from three previously published protein or RNA human atlases [8,9] to augment our proteome distribution atlas to generate a comprehensive baseline map of protein/transcript abundance across 29 different human tissues and blood cells. We demonstrate that the surrounding tissues specifically contribute to the plasma proteome and that the composition of tissue-enriched proteins change in disease states such as myocardial injury, pancreatitis, and infections.

## Results

### Construction of a blood-plasma centered tissue and cell distribution atlas

To investigate if in-depth analysis of tissue and cell proteomes can be used to infer the origin of proteins detectable in plasma, we collected 18 healthy tissues and isolated 8 cell types from three or more individuals per organ/cell type (**Fig. 1a**). The organs and cells were selected based on the likelihood of impacting the plasma proteome and included all major vascularized organs and the most abundant cells-types in blood. The tissue biopsies were collected from healthy tissues of focal disease, healthy volunteers or during prophylactic or reconstructive surgery (see extended method section for more details). The cells were isolated from blood samples from healthy volunteers using cell-specific isolation protocols. The cell purity was determined using flow cytometry (**Sup. Fig1**).

**Figure 1:**
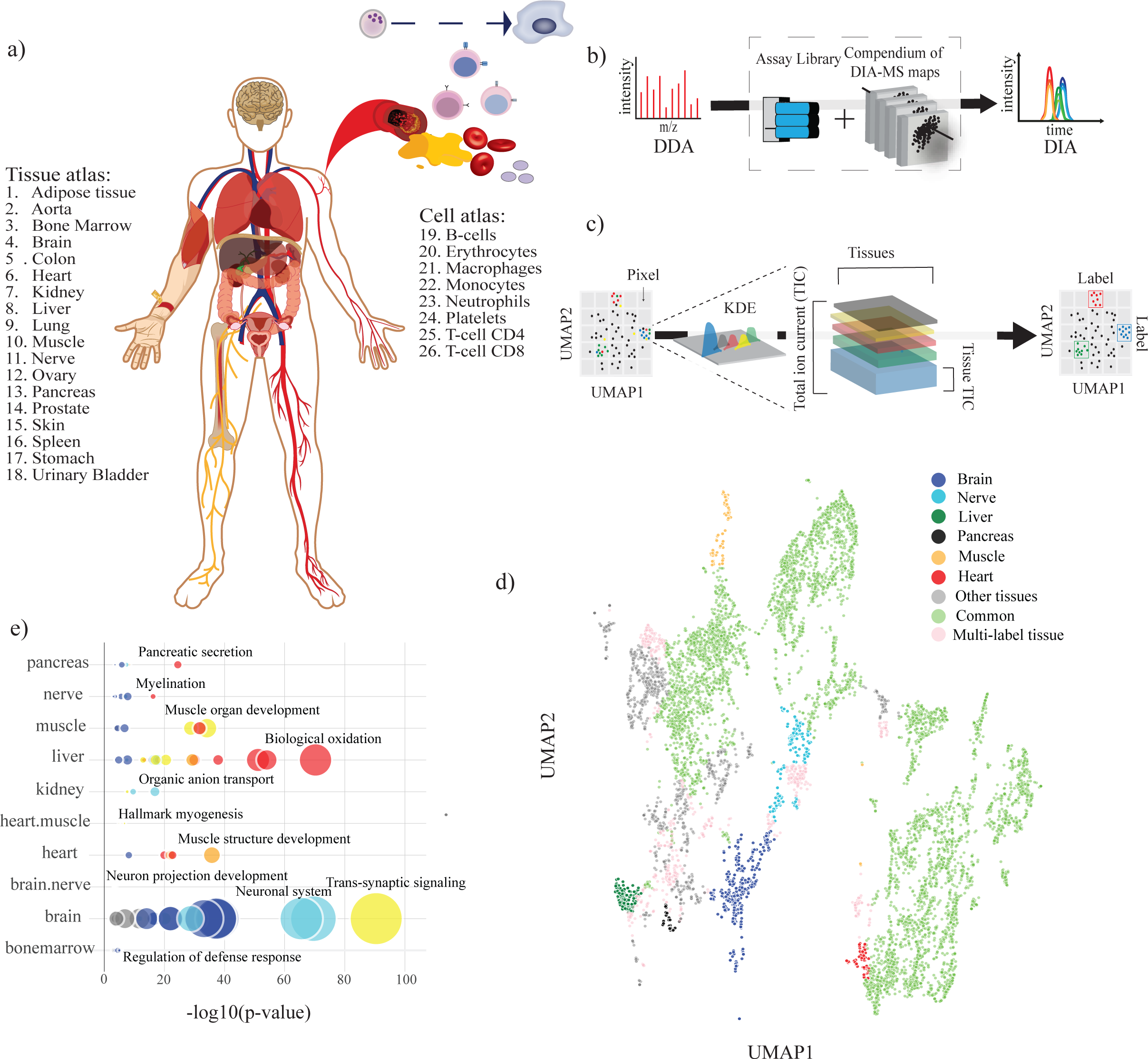
Outline of samples and experimental strategy. (A) Organs and cells predicted to have a high likelihood of impacting the blood plasma proteome were collected or isolated from at least three individuals per organ/cell type (B) A schematic overview of the mass spectrometry experimental design. The tissues and cells were homogenized and fractionated by SDS-PAGE, followed by LC-MS/MS analysis in DDA-mode to construct a spectral assay library. Unfractionated tissue and cell samples were reanalyzed using DIA-MS to generate a compendium of 156 proteome-maps and the spectral library was used to extract protein identities and quantities (referred to as HATLAS). (C) Schematic overview of tissue– or cell-label assignment strategy. The dimensionality of the dataset was reduced using UMAP to group proteins with similar tissue or cell abundance profiles across the analyzed tissue. We then applied a Gaussian KDE for each protein, using each tissue’s relative fraction of the total ion current. Each protein-tissue pair was then projected onto a 400×400 pixel image, representing the tissues as a channel, summing the contribution for each neighbor protein. The contribution for each protein was summed for each pixel. We classified each pixel by removing low-abundant channels and assigning it to a single label if one of the remaining tissues contributed more than 50% of the total signal. Each pixel was then classified as either belonging to a single or to multiple tissues or designated as a common protein. Last, we identified the closest pixels to each protein and used the labels for the pixel to label the protein. (D) Resulting UMAP of all identified proteins in the proteome atlas, with selected single– or multilabel highlighted. Functional enrichment analysis was performed for all identified tissues and the results for selected single– and multilabels are shown (E). The color represents enrichment; red, orange, yellow, cyan, blue and gray represent 5, 4, 3, 2, 1, fold enrichment, respectively and the glyph size represents the number of observed successes, e.g., the number of proteins in the cluster of that annotation.

All samples were prepared for mass spectrometry (MS) analysis followed by off-line fractionation and the fractions were analyzed using data-dependent acquisition mass-spectrometry (DDA-MS). The resulting DDA-MS data was converted to a spectral library covering 10786 proteins. For quantification, unfractionated tissue and cell samples were reanalyzed using data-independent analysis mass spectrometry (DIA-MS) and quantified using the spectral library to assess protein abundance (**Fig. 1b**). In total, 9960 unique proteins were quantified at a 1 % false discovery rate. The data was used to separately construct a blood cell-based and a tissue-based atlas to enable inference of both tissue and cell-based protein origins. Over 90.5 % of all identified proteins were shared between the atlases, demonstrating a considerable overlap between the tissue– and cell-based proteomes **(Sup. Table 1a and 1b).**

Previous studies have observed that most proteins are produced in several tissues, in varying quantities, making it challenging to categorize protein-tissue associations. ([1,10]). To classify protein-tissue associations more objectively, we first used uniform manifold approximation and projection (UMAP) [11] of the 9960 normalized protein intensities across the analyzed tissue proteomes. In the next step, a weighted Gaussian kernel density estimation (KDE) was applied to each protein for each tissue (**Fig. 1c**), using each tissue’s relative fraction of the total ion current, to create a multi-channel image with an x and y resolution of 400×400, one for each tissue. The contribution for each protein was summed for each pixel. We classified each pixel by removing low-abundant channels and assigning it to a single label if one of the remaining tissues contributed more than 50% of the total signal. Each pixel was then classified as either belonging to a single or to multiple tissues or designated as a common protein. The label a pixel received was then propagated to the proteins that fell within that pixel. If a protein was assigned to two or three tissue labels, they were denoted as multilabel proteins, and proteins assigned to four or more tissue labels were denoted as common. The same strategy was applied for blood cell proteome maps.

In this way, proteins with similar abundance profiles across the analyzed tissues were grouped together and assigned to a corresponding tissue/cell label (**Fig. 1d)**. The proteins were categorized into three main groups: i) proteins that were enriched in one tissue– or cell-type (tissue– or cell-enriched proteins, n=2317 and n=5289 respectively), ii) proteins enriched in more than one tissue/cell (multilabel tissue or cell proteins, n=782 and n=1847) and iii) proteins present in similar amounts in all tissues/cell-types (common tissue or cell proteins, n=6822 and n=1920). To estimate the accuracy of the label assignments, we performed functional enrichment analysis [11]. Most organ/cell labels generated biological functions matching the function of the respective organ, such as contractile proteins enriched in heart or muscle, drug metabolism in the liver (cytochrome P450 enzymes) and protein associated synaptic signaling in the brain (**Fig. 1e**, **Sup. Table 1e**). In addition, the strategy enabled us to identify several tissue-enriched proteins with known tissue association, such as cardiac troponins, pancreatic amylase, and pulmonary surfactant-associated proteins **(Sup.** Fig 2**, Sup. Table 1a and 1b).** The proteins assigned to the multilabel category represent a group of proteins that are shared between two or several organs. The class of multilabel proteins was most frequent between tissues with similar functions, such as heart/muscle and brain/nerve. These results demonstrate that the compendium DIA-MS tissue maps and the UMAP/KDE classification strategy generated an unbiased definition of tissue protein labels between one or several tissues, expanding the number of proteins that can be monitored during pathological conditions.

### A global distribution atlas of the human proteome

To create a broader consensus map of the protein distribution across multiple atlases, we downloaded the raw data from previously published RNA/protein atlases and integrated them with our tissue– and cell proteome atlases. The combined atlases consisted of four tissue atlases (two protein and two RNA) and two cell atlases (one protein and one RNA), containing a total of 18383 transcripts and 11976 proteins across 21 different tissues and eight blood cells ([8,9]). To objectively assign tissue labels to all the atlases, we reanalyzed the downloaded data with the same UMAP/KDE classification strategy described above **(Sup. Table 1a-b and 2a-d).** Every RNA/protein analyte received a weighted score from each atlas based on the underlying label assignment (single label, multilabel or common) and how often it was identified in the different atlases. The weighted score generated from the individual atlases was summed up to obtain a global label score (GLS), and the tissue or cell label with the highest GLS was used as the primary annotation (see extended method section for more details) **(Fig. 2a**). The GLS ranged from 0-4 and reflects the confidence of a given protein-tissue label assignment.

**Figure 2:**
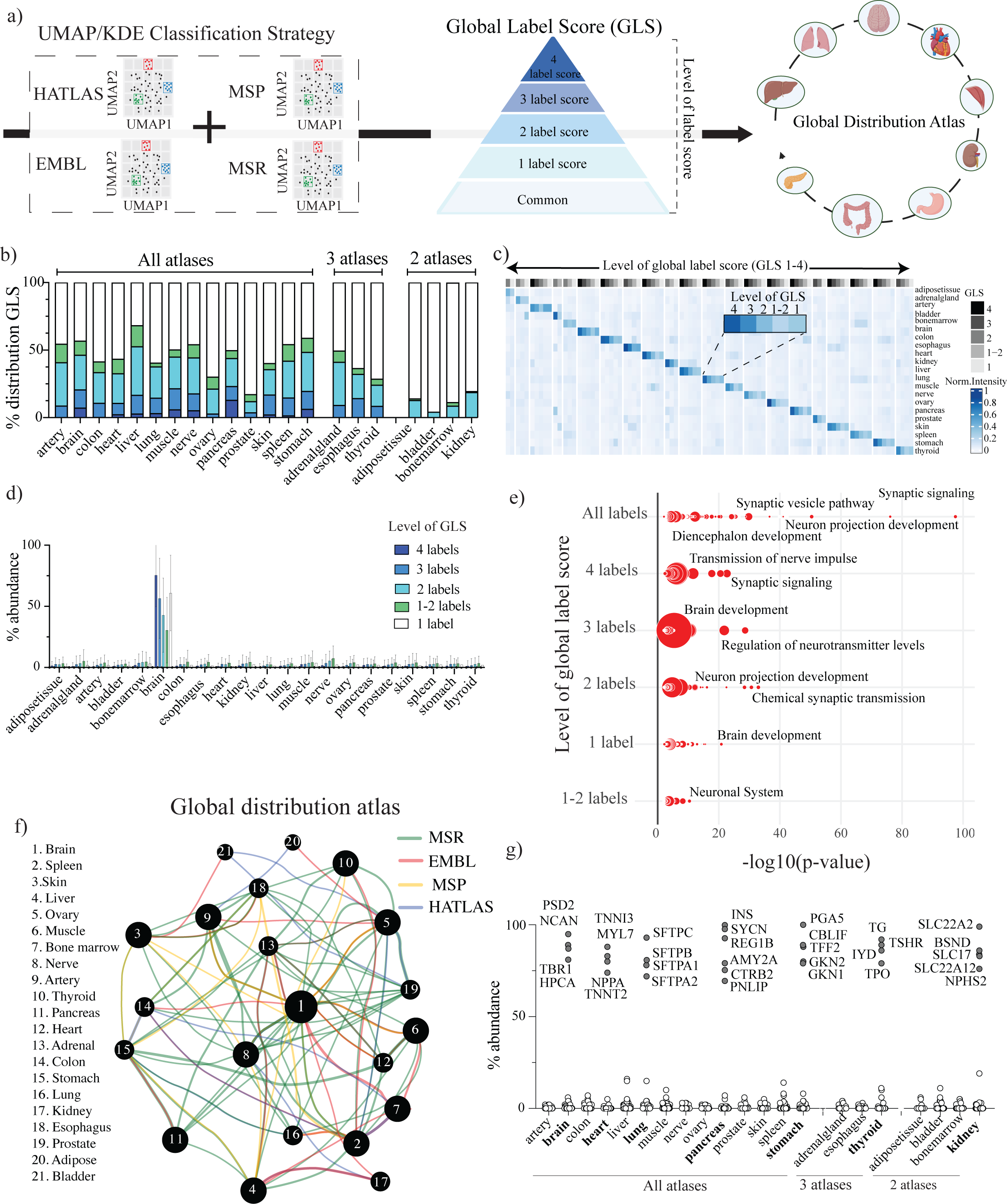
A Global distribution atlas of the human proteome. (A) The KDE on UMAP projections were used to assign tissue-protein/RNA labels for each atlas (referred to HATLAS, EMBL, MSP and MSR). Each RNA/protein received a weighted score based on the tissue labels assigned from the respective atlas. The weighted score for each protein was summed up to obtain a global label score (GLS). The tissue or cell label with the highest GLS was used as primary annotation, thus using the label score to integrate the individual atlases into a global distribution atlas. (B) Bar graph displaying the level of global label score for each tissue. (C) Heatmap depicting the average protein abundance within each level of GLS for tissue-enriched proteins. (D) The distribution of normalized protein intensities for brain-enriched proteins for each level of the global label score. The bars represent the average abundance across each atlas and error bars indicate standard deviation. (E) Functional enrichment analysis of transcripts/proteins with different levels of GLS exemplified by brain-enriched proteins. The size of the nodes represents the number of assigned tissue proteins. (F) An integrated network of protein tissue assignments based on the global distribution atlas. The nodes represent assigned tissue-enriched proteins, and their size reflect the number of proteins within each tissue. The edges and the different colors represent appointed multilabel proteins from the individual atlases and are color-coded accordingly. (G) Examples of proteins with high tissue specificity across the global distribution proteome atlas.

Protein assignments with a GLS of 4 highlight proteins that were confidently labelled to a single tissue by all atlases. These high-confident proteins were relatively few, accounting for around 10% of the total tissue assignments. Over 90% of the proteins that were assigned to a given tissue were missed by one or more of the atlases most likely due to biological or experimental variation **(Fig. 2b**). Most of the time, proteins/transcripts with a low GLS were only identified in a one or two atlases. To investigate the confidence of the proteins with a global label score below 4, we quantified the average abundance within the different levels of global label scores of 1, 1-2, 2, 3 and 4 and how the intensities for these proteins/transcripts were distributed across the analyzed tissues **(Fig. 2c**). This analysis showed that even RNA/proteins with lower GLS (<2) are still predominantly found in the assigned tissue, as further exemplified by the quantitative abundance distribution for brain-enriched proteins in **Figure 2d** and **Sup.** Fig 3. We also performed functional enrichment analysis of the protein/transcripts within each level of the global label score, illustrated in the group brain-enriched proteins (**Fig. 2e**). Notably, proteins with low GLS were typically enriched for tissue-specific functions and there was an extensive overlap with the enrichment results observed in protein groups with high GLS (**Sup. Table 2e-f**). These results indicate that the abundance of tissue-enriched proteins ranges from being specifically produced in one tissue type to more modestly enriched proteins that are more challenging to pick out by one experimental approach using one tissue sample from one individual.

To visualize the tissue assignments, we plotted the global distribution atlas in a network graph. The edges represent multilabel proteins from each atlas, and each node depicts tissue proteins defined by the GLS (**Fig. 2f and Sup. Table 2g**). On average, tissues were assigned 261 proteins whereas the brain and spleen had the highest number with 1012 and 475 proteins respectively. Figure 2g shows the normalized average expression of selected tissue-specific proteins **(Fig. 2g and Sup. Table 2h)**. Among the cell types, erythrocytes (n=959), platelets (n=737) and neutrophils (n=691) had the highest number of cell-enriched proteins whereas heart/muscle, bone marrow/spleen, and brain/nerve were among the most prevalent multilabel protein groups.

Collectively, these results demonstrate that each atlas provides unique information. The strategy to classify protein-tissue/cell association and to combine multiple atlases objectively creates a broader and more robust definition of the molecular distribution across tissues and cells.

### Data-driven definition of primary plasma proteins

Proteins with primary functions in plasma, such as coagulation and complement proteins, are predominantly produced in the liver. For this group of proteins, referred here to as primary plasma proteins, it can be expected that RNA levels are high in the liver, but the protein levels are low since these proteins are actively secreted into the plasma upon protein translation. Our global distribution atlas, which contains RNA and protein abundance data, provides an opportunity to define the group of primary plasma proteins in a data-driven manner. Using the global distribution atlas, we selected proteins with liver labels in the RNA atlases and removed proteins with assigned liver labels in the protein atlases. For the selected proteins, we determined the relative abundance ratio in the liver, contrasted the other tissues in each atlas and compared the results between the RNA and protein atlases. The strategy allowed us to identify proteins with 5-10,000 higher liver abundance ratio in the RNA atlases compared to the protein ones. Of these, 114 proteins were quantified with at least two peptides in plasma from a cohort of healthy individuals[12]. (**Fig. 3a**). Functional enrichment analysis revealed that these primary plasma proteins were involved in the principal functions of blood plasma, such as the complement and coagulation cascades (**Fig. 3b**). Typical proteins among these primary plasma proteins were coagulation factors, complement proteins, albumin, apolipoproteins, and acute-phase proteins **(Sup. Table 3a).**

**Figure 3:**
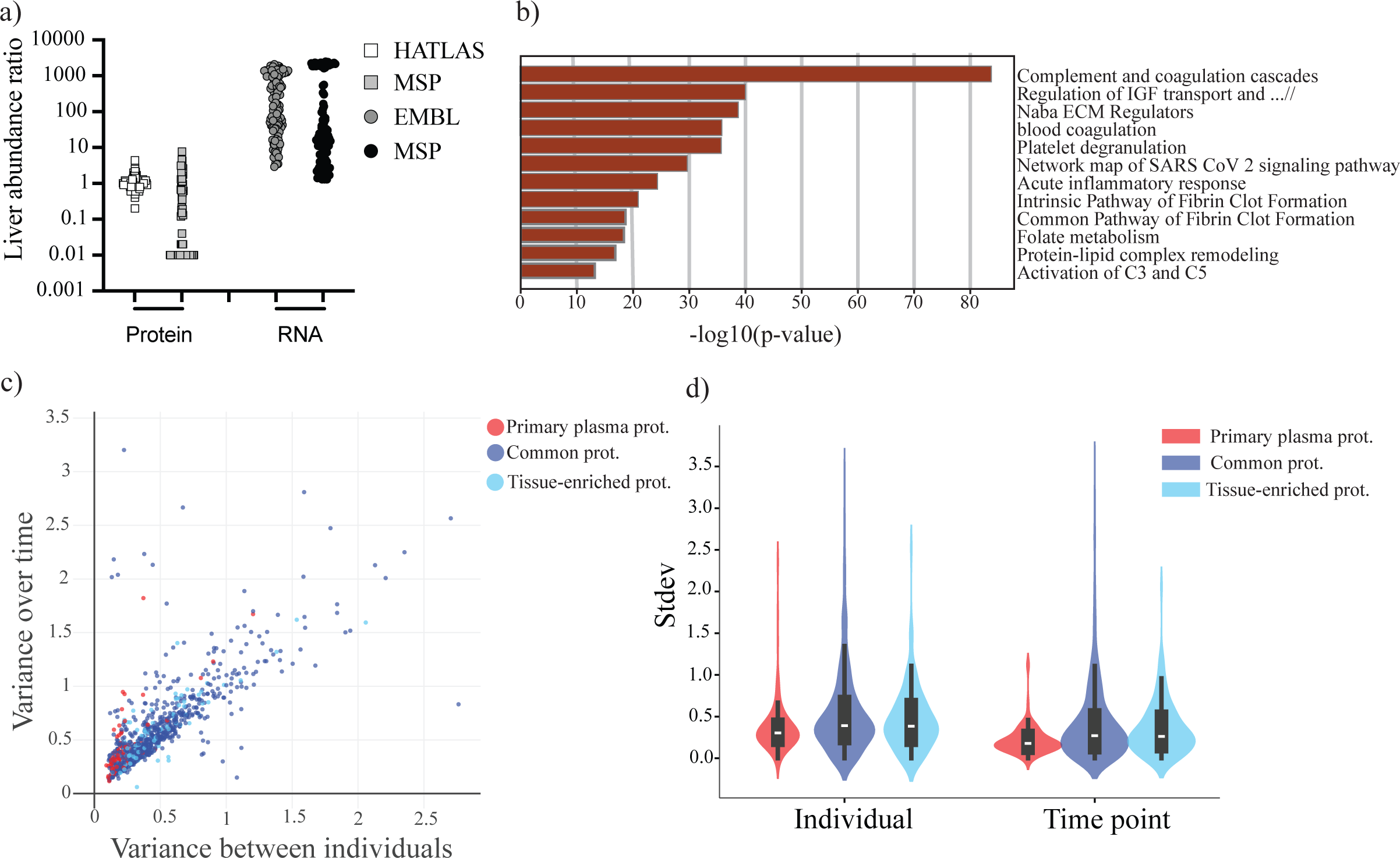
A data-driven strategy to define primary plasma proteins. A data-driven strategy comparing differences in abundance between RNA and protein levels in the liver to define primary plasma proteins. (A) The dot plot displays 114 proteins with at least a 5-fold higher liver abundance ratio in RNA atlases compared to protein atlases. (B) The horizontal bar graph shows the functional enrichment analysis of the defined primary plasma proteins. Five plasma samples from 10 healthy volunteers collected over five weeks were analyzed using DIA-MS. Protein identities and intensities were extracted from the compendium of DIA-MS maps using the human proteome distribution spectral library. (C) Scatter– and (D) violin plots displaying the calculated variance of the quantified tissue-enriched, common tissue– and primary plasma proteins for all individuals and all the different time points. Red denotes primary plasma proteins. Light dots denote tissue-enriched proteins, and dark blue denotes proteins defined as common.

Since primary plasma proteins are actively secreted into the plasma and not the result of cell necrosis, we speculated that they are under more strict homeostatic control than tissue-enriched proteins. To test this, we used time-resolved blood plasma samples from healthy volunteers mentioned above. In this cohort, five samples were taken from 10 individuals on a weekly basis over five weeks and analyzed using DIA-MS. We calculated the variance of quantified tissue-enriched proteins, commonly shared tissue proteins and primary plasma proteins for all individuals and all the different time points. This analysis revealed that variation over time for different individuals was marked higher than the variation within one individual (**Fig 3c-d**). Furthermore, the variance of protein levels for primary plasma proteins was lower compared to both commonly shared and tissue-enriched proteins, indicating a stricter control of this protein group. Collectively, the data-driven definition of primary plasma proteins allows improved monitoring of tissue-derived proteins in plasma. Further, these results demonstrate that several tissue-enriched proteins detectable in plasma have a higher interindividual variation than primary plasma proteins under normal physiological conditions.

### Pathological changes of tissue-specific protein signatures in plasma

The defined tissue-enriched proteins and the data-driven definition of primary plasma proteins provide new possibilities to monitor the dynamics of the plasma proteome under pathological conditions. To test this, we collected blood plasma samples from three different patient cohorts at the emergency department (ED) as a proof of concept. The cohorts consist of patients with pancreatitis (n=14+10), myocardial injury (MI) (n=25+25), and infections with different microbial etiology (n=47+40) along with their respective controls (**Fig. 4a-c**). The three conditions have different underlying pathophysiology and well-established clinical biomarkers such as troponin T, amylase, and c-reactive protein (CRP). The plasma samples were analyzed using DIA-MS and the tissue atlas spectral library was used to extract protein intensities to generate a compendium of 161 DIA-MS proteome maps **(Sup. Table 4a-c).**

**Figure 4:**
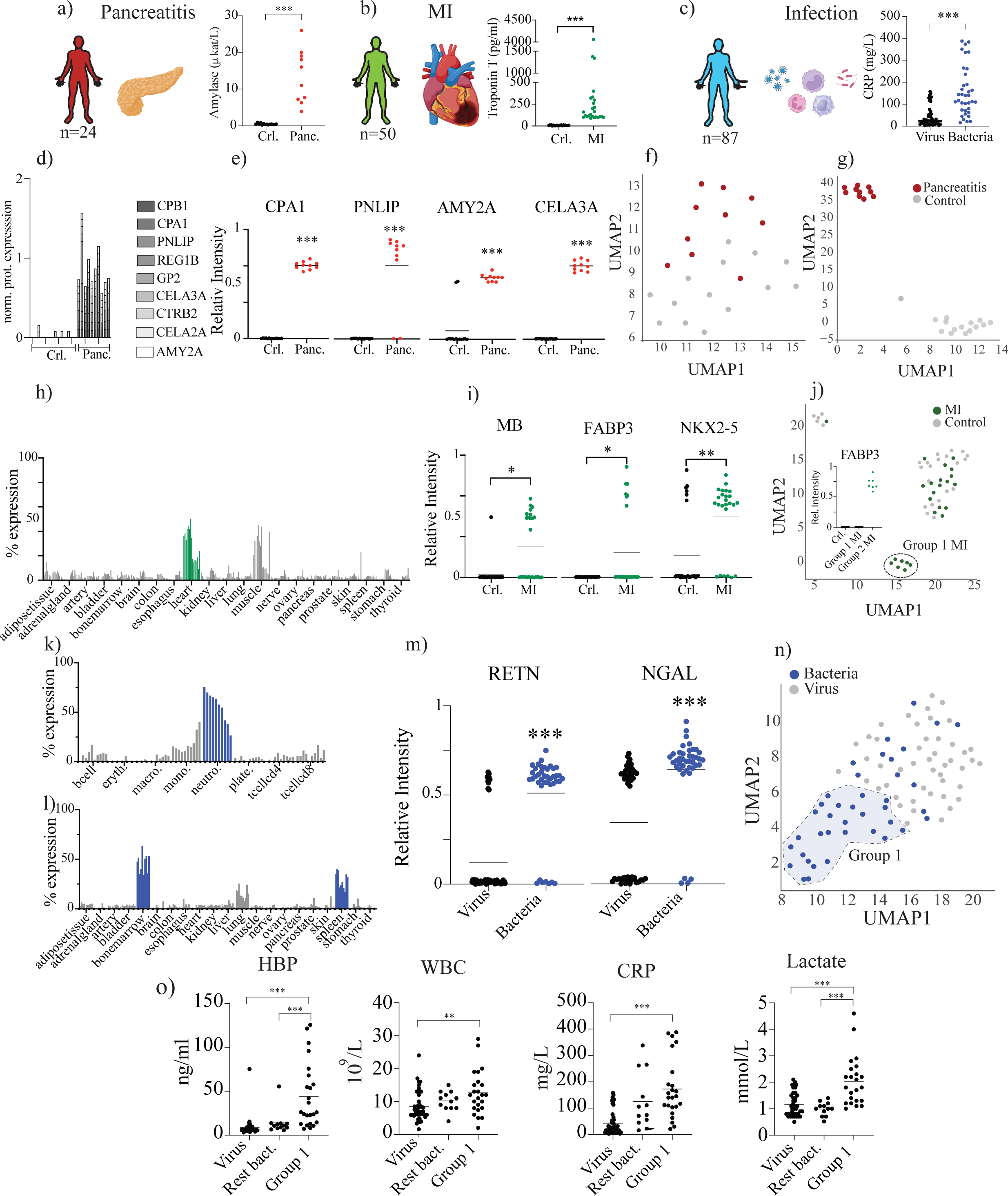
Pathological changes of tissue protein signatures in plasma. (A-C) Overview of the pancreatitis, MI and infection cohorts enrolled at the emergency department with their associated clinical biomarkers (Amylase, Troponin T and CRP). The blood plasma samples from all patients were analyzed using DIA-MS and the protein intensity was extracted by spectral library to generate a compendium of 161 DIA-MS proteome maps. The origin of all proteins was inferred using the global distribution atlas. (D) The bar graph depicts the normalized protein abundance of nine pancreas-enriched proteins in the pancreatitis plasma cohort (control vs pancreatitis). (E) Dot plot illustrating the abundance pattern of four pancreas-enriched proteins that were significantly elevated in pancreatitis plasma relative to healthy controls. (F-G) Uniform Manifold Approximation and Projections (UMAP) of the pancreatitis cohort using all identified plasma proteins or filtered only using pancreas-enriched proteins defined by the global distribution atlas. (H) Bar graph showing the average abundance level of the 15 identified heart-enriched proteins across the different tissues. (I) Dot plot illustrating the abundance pattern of three heart-enriched proteins that were significantly elevated in MI plasma relative to control plasma (J) UMAP on individual patients in the MI cohort, filtered on heart-enriched proteins and the dot plot shows the abundance pattern of the protein FABP3 in a subgroup (Group 1 MI) of patients identified in the UMAP. (K-L) Average abundance of nine neutrophil-enriched proteins were identified in plasma from the infection cohort across the different cells and tissues respectively. (M) Dot plots illustrate two neutrophil proteins significantly elevated in the plasma from patients with bacterial infection relative to viral infection. (N) UMAP on individual patients in the infection cohort, filtered on neutrophil-, spleen– and bone marrow-enriched proteins, identifying a subgroup of patients (Group 1). (O) Levels of white blood cell count, CPR, lactate and heparin-binding proteins in Group 1, remaining bacterial infection and viral infection. Mann-Whitney tests were performed for the indicated comparisons. *p < 0.05; **p < 0.01, ***p < 0.001.

In total, we identified 17 pancreas-enriched proteins in the plasma samples from the pancreatitis cohort. The normalized abundance pattern of these proteins across the different tissues are shown in **Supplementary figure 4**. Nine of these proteins had a GLS of 4, six had a score of 3, and two had a score of 2. Further, 8 of these proteins, including pancreatic alpha-amylase, were significantly elevated in plasma from pancreatitis patients relative healthy controls (**Fig. 4d-e**). Separating the patient cohort based on all identified plasma proteins showed that the entire plasma proteome could not separate the pancreatitis patients from their respective controls (**Fig. 4f**). However, data-driven filtering of the plasma proteome, selecting only pancreas-enriched proteins defined in the global distribution atlas, markedly improved the separation between pancreatitis patients and the controls (**Fig. 4g**).

The second cohort was composed of patients with chest pain admitted to the ED, with and without myocardial injury (MI). The MI group had elevated Troponin T levels compared to baseline (**Fig. 4b**). In this cohort, the global distribution atlas enabled the identification of 15 heart-enriched proteins in plasma, and their average abundance pattern from all the atlases across the different tissues is shown in **Fig. 4h**. Three proteins, myoglobin (MB), Fatty acid-binding protein, heart (FAB3P) and Homeobox protein Nkx-2.5 (NKX2-5), were significantly elevated in plasma from patients with myocardial injury relative to control (**Fig. 4i**). Interestingly, MB and FAB3P have already been proposed as cardiac biomarkers [13], whereas NKX2-5 acts as a transcription factor essential for heart development[14]. Based on the levels of FABP3, we identified a subgroup of patients with myocardial injury that clustered, indicating that the release of heart-enriched proteins is not uniform in this cohort (**Fig. 4j**).

In the final patient cohort, we included patients with infection caused by different microbial agents (n= 47 viral infection, n= 40 bacterial infection). Neutrophils are key players in the innate immune system and essential in orchestrating the early host response against invading pathogens. To target neutrophil-enriched proteins [15], we combined the tissue– and cell atlases since individual cells reside within different organs/tissues and there is an extensive protein overlap between the two atlases. For this purpose, we performed data-driven filtering of the plasma proteome using cell-enriched neutrophil proteins and tissue-enriched spleen and bone marrow proteins as these organs are rich in neutrophils. The abundance level for a subset of these RNA/proteins from all the atlases across different cell types and organs confirms that the proteins are abundant in the selected cell and tissues (**Fig. 4k-l**). Nine neutrophil-derived proteins were significantly elevated in plasma from patients with bacterial infection compared to viral infections. (see examples in **Fig. 4m**). Akin to the pancreatitis and MI cohort, filtering the plasma proteome on the neutrophil-specific proteins separated 67.5 % of the patient’s bacterial infections, referred to as group 1 (**Fig. 4n**). This subgroup of patients with bacterial infection had higher white blood cell count and CRP levels than the group with viral infection. Further, group 1 also has higher heparin-binding protein (HBP) and lactate levels than the remaining patients with bacterial infection, suggesting a more pronounced hemodynamic instability within the subgroup (**Fig. 4o**) [16,17].

### Comparison of pathological plasma proteome changes across different patient cohorts

To test if changes in tissue-enriched protein signatures are specifically associated with a given pathophysiological mechanism or a specific condition., we compared tissue– and cell-enriched protein variations across the three patient cohorts. The data showed increasing levels of acute phase proteins in all three diseases (**Fig. 5a**). These results were anticipated since it has previously been described that acute phase proteins are mobilized during acute coronary syndrome [13] and that the pathophysiology of sepsis and pancreatitis is associated with systemic inflammation. In contrast, changes in the expression of pancreas-enriched proteins in plasma was specifically associated with pancreatitis (**Fig. 5b**). Further, the levels of the heart-enriched proteins FAB3P and NKX2-5 were, to a greater extent, elevated in patients with myocardial injury compared to the other cohorts. Myoglobin, on the other hand, was significantly elevated in patients with bacterial infection (**Fig. 5c**). Sepsis is associated with cardiac dysfunction, but the underlying pathophysiology differs compared to acute coronary syndrome [18], which may explain these observations. Finally, neutrophil-enriched proteins increased more in bacterial infection, although a partial overlap was observed in patients with myocardial injury (**Fig. 5d**). Thus, the results showed both common and unique changes in the tissue-enriched protein abundance levels across all patient cohorts.

**Figure 5:**
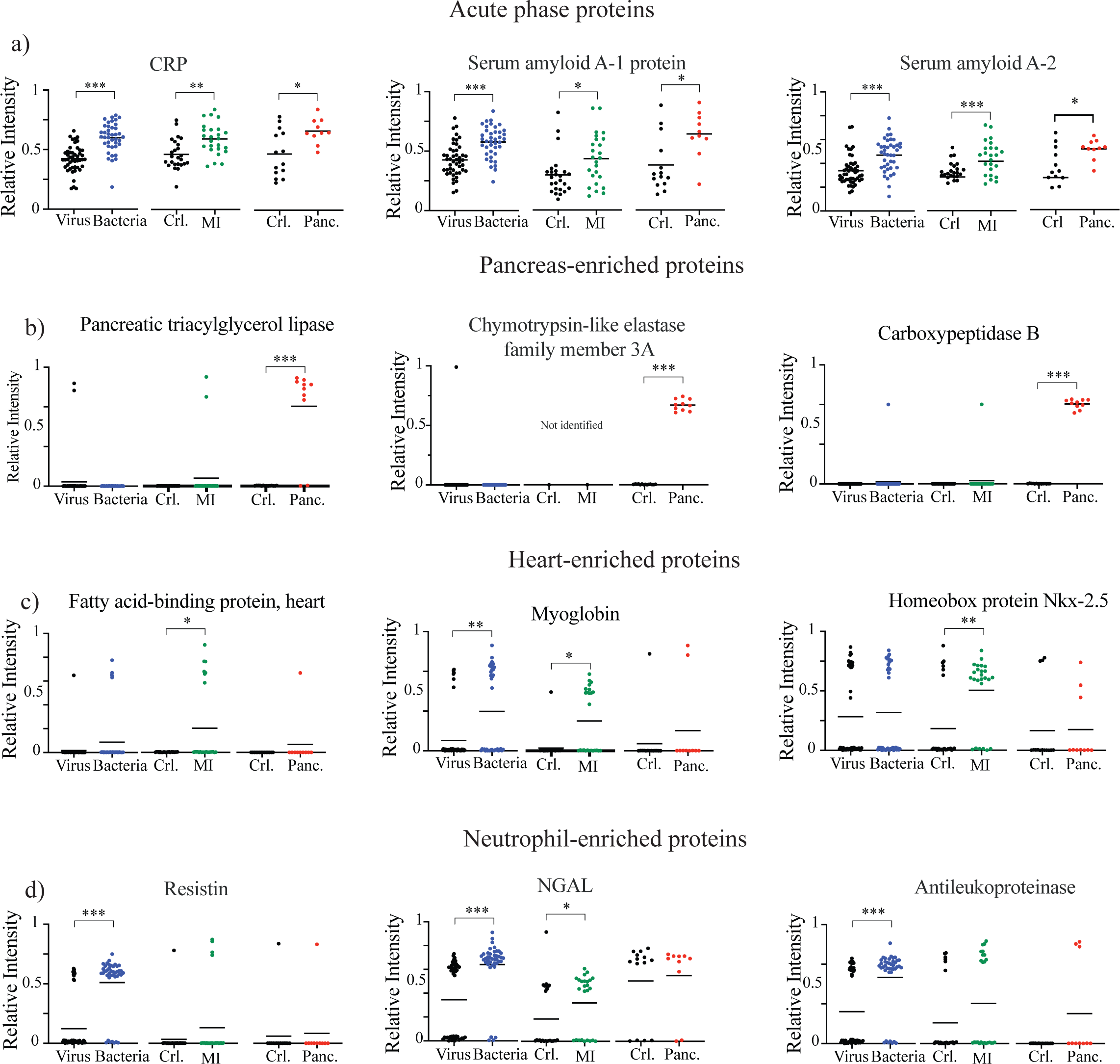
Comparison of pathological plasma proteome changes across different patient cohorts. The Global distribution atlas combined with a compendium of 161 DIA-MS proteome maps were used to compare changes of tissue-enriched– and primary plasma protein signatures across the patient cohorts. (A-D) Dot plots exemplify the abundance pattern of acute phase-, pancreas-enriched-, heart-enriched and neutrophil-enriched proteins in plasma across the three patient cohorts. Mann-Whitney tests were performed for the indicated comparisons. *p < 0.05; **p < 0.01, ***p < 0.001.

In conclusion, the results show that surrounding tissues and cells alter the plasma proteome composition during disease, which can be used to reveal disease-specific protein patterns not found using global plasma proteome analysis. These results further demonstrate that the global distribution atlas connected to DIA-MS enables reproducible quantification of tissue-specific proteins with inferred tissue origin directly in clinical plasma samples.

## Discussion

The plasma proteome has over the years been extensively investigated as a source for understanding diseases and for biomarker discovery studies. However, the incomplete understanding of how surrounding tissues and cells impact plasma proteome composition complicates such investigations.

In this work we constructed a proteome distribution atlas to infer the most likely protein origin for all proteins detectable in plasma to facilitate the identification of tissue-specific protein signatures in plasma. To create a broader consensus map of the protein distribution across multiple atlases, the proteome atlas was interfaced previous large-scale cataloguing effects to reduce variation from measurement technology to assemble a broader global distribution score across several tissues and cells. Interestingly, this analysis revealed that only 10 % of the tissue-enriched proteins had the highest global label score, showing the importance of using multiple atlases. Many factors, such as the underlying cell composition within a biopsy and individual and technical differences, influence tissue RNA/protein abundance levels.

We predict that a combination of the global distribution atlas and large-scale DIA-MS plasma analysis can be extended to measure how tissue proteins are regulated in many other pathological conditions, improving our understanding of the processes that influence plasma proteomes during disease. In this regard, kernel density estimation to define tissue-protein assignments is beneficial as it offers a flexible, data-driven solution that can be adjusted based on experimental question rather than relying on manual rule-based alternatives [19] The resource presented here provides a starting point to further increase our knowledge of how different pathophysiological processes distort the protein composition of healthy plasma to explore the underlying pathophysiological mechanisms.

## Material and methods

### Organ collection and preparation

The collection of human samples relies on collaborative work between preclinical researchers, mass spectrometry experts and surgeons at the Departments of General Surgery, Cardio-thoracic-, Neuro-, Hand Surgery, Gynecology and Urology, at Skånes University Hospital and the hospital of Helsingborg. Twenty-six tissues and cell types were collected with approved ethical permissions (see below) and biobank permission from the local ethical committees. Three knife biopsies per tissue approximately 4×4 mm (minimum of 50 mg; or for the peripheral nerve at least 20 mm long due to its smaller diameter) were collected during surgery from different patients from every organ. The tissues were immediately put on ice and stored at – 80°C within one hour of collection. The samples have been isolated from the following surgical procedures:

### Pancreas

Pancreatic tissue was retrieved from a surgical procedure called pancreaticoduodenectomy, including resection of the duodenum, the head of the pancreas, distal bile ducts and the gall bladder. The operation was performed due to malignancy of the pancreas. Healthy pancreatic tissue was retrieved from the non-tumorous tissue in the specimen (DNR 2014/764 and DNR2015/668).

### Liver

Liver tissue was collected from one patient who underwent a benign liver cyst resection and one patient who underwent pancreaticoduodenectomy (see description above). A third sample was collected from a patient who underwent a local resection of colorectal metastasis (no prior chemotherapy treatment), and healthy tissue from a different part of the liver was taken (DNR 2014/764 and DNR2015/668).

### Spleen

Spleen tissue was retrieved from normal spleens on two patients who underwent distal pancreatic resection including splenectomy for premalignant pancreatic diagnosis. A tissue sample from a spleen harvested during organ procurement was acquired (DNR 2014/764 and DNR2015/668).

### Heart, aortic vessel, adipose tissue and muscle tissue

Epimyocardial and intermediate size vessel biopsies were collected during open heart surgery (coronary artery bypass grafting). Tissue samples were harvested from the right atrial auriculum, the left internal mammary artery and from the ascending aortic wall during anastomosis of the proximal vein used as a bypass graft in the procedure. During the same procedure also adipose and muscle tissue was collected (DNR 2014/764 and DNR2015/668).

### Stomach/ ventricle and colon

Biopsies were taken from patients with macroscopic normal upper (gastroscopy) or lower (colonoscopy) endoscopy (DNR 2014/764 and DNR2015/668).

### Ovaries

Ovaries were surgically removed from patients with hereditary accumulation of breast and ovarian cancer but without BRCA mutation (EPN 2004/558).

### Skin

A 5 mm cutaneous punch including dermis, and subcutaneous fat was taken from healthy volunteers (DNR 2014/764 and DNR2015/668)..

### Prostate and urinary bladder

Tissue samples from the prostate and urinary bladder were obtained from four patients undergoing TURP (TransUrethral Resection of the Prostate). From the bladder samples were taken from both the side wall and the trigonal area representing both endodermal and mesodermal origin. Three of the four patients had a known prostate cancer, but pathologic examination did not indicate malignancy in the removed specimen, so the prostate samples represent benign prostate enlargement. The tissue from the urinary bladder was normal (Dnr 790/2005 2006-01-17).

### Kidney

Patients included were operated with radical nephrectomy, involving removal of the entire kidney. The procedure was accomplished using either robot assisted approach or laparoscopic surgery. The patients were operated due to benign oncocytoma, papillary kidney cancer or clear cell kidney cancer. The healthy kidney tissue was perceived from the opposite pole of the cause of the operation. The kidney was microscopically investigated to make sure the healthy tissue was without any involvement of disease (DNR 2014/764 and DNR2015/668).

### Lung

Healthy lung tissues were collected from organ donors, with no prior history of lung disease according to their medical register and this was further confirmed by interviews with their closest relatives. This study was fully approved by the Swedish Research Ethical Committee in Lund (FEK 91/2006) and in Gothenburg (FEK675–12/2012). From distal lung specimens, samples were taken from locations just below the pleural surface as previously described[20].

### Brain

Brain tissue was collected from otherwise healthy patients who underwent neurosurgery due to epilepsy (Dnr 212/2007). The surgery was performed as a temporal lobe resection including hippocampus and parts of the amygdala.

### Peripheral nerve

Up to 30 mm long nerve biopsies were taken (remaining part after procedure; free from fat) from the sural nerve at lower leg in connection with a procedure to use the sural nerve in reconstruction of another lacerated nerve in otherwise healthy patients (Dnr 2016/5 LU, Dnr 2016/98, Dnr 2024-01893-0).

### Neutrophil preparation

Whole citrated human blood from healthy donors was layered on Polymorphprep (Axis-Shield) and centrifuged at 400 × g for 35 min at 20°C according to manufactures instructions. The neutrophil layer was recovered and suspended in 50 mL calcium– and magnesium-containing PBS. After centrifugation at 350 × g for 10 min, erythrocytes were removed by hypotonic lysis for 20 seconds. The cells were then pelleted at 250 × g for 5 minutes at 20°C and resuspended in Na medium (containing 5.6 mM glucose, 127 mM NaCl, 10.8 mM KCl, 2.4 mM KH2PO4, 1.6 mM MgSO4, 10 mM Hepes and 1.8 mM CaCl2; pH adjusted to 7.3 with NaOH). Cells were further purified with Fluorescence-activated cell sorting (FACS) (see method section Flow Cytometry and Antibodies) (DNR 657/2008).

### Erythrocytes

Whole citrated human blood from healthy donors was layered on Polymorphprep (Axis-Shield) and centrifuged at 400 × g for 35 min at 20°C. The bottom erythrocyte layer was collected and washed three washing times (400 x g, 10 minutes). The cells were resuspended in PBS and the cell purity was verified using flow cytometry (DNR 657/2008).

### Platelets

Whole citrated human blood from healthy donors was centrifuged at 160 x g for 10 minutes at 20°C to obtain platelet rich plasma (PRP). The PRP was collected followed by a new centrifugation step (160 x g, 10 minutes, 20°C). The PRP was recovered again, and the cells were pelleted at 800 x g for 10 minutes at 20°C and resuspended in PBS. This washing step was repeated once. The cell purity was verified using flow cytometry as described below (DNR 657/2008).

### Lymphocytes and monocytes

Mononuclear cells were isolated from a leukocyte unit (obtained from Reveos automated blood processing system) using Lymphoprep density centrifugation as described by the manufacturer (Axis-Shield). The collected mononuclear cell layer was washed twice (400 x g, 10 minutes, 4°C) and resuspended in PBS. Cells were further isolated into B-cells, Monocytes, CD4 T-cells and CD8 T-cells using FACS (DNR2014/764).

### Macrophages preparation

Human monocytes were isolated from a leukocyte unit (obtained from Reveos automated blood processing system) by Lymphoprep density centrifugation as per the manufacturer’s instructions (Axis-Shield), followed by an anti-CD14-coated microbead-based purification step. Purified CD14+ monocytes were seeded at a density of 1 million cells/well in 6-well plates in RPMI 1640 (Life technologies) with 10% serum and 20μg/ml recombinant human MCSF (Peprotech) and cultured for 8 days. Cells were taken in PBS and prepared for mass spectrometry. Macrophage purity was determined to be 85 >% using immunofluorescence with macrophage markers anti-CD68 (5μg/ml) and –CD163 (2μg/ml) (Biolegend) (DNR2014/764).

### Bone marrow

Bone marrow aspirates from healthy donors were collected at the Department of Hematology at Skåne University Hospital in Lund, Sweden (Dnr 2021-04046). The Bone marrow donors was diluted to a ratio 1:1 with PBS and layered on Ficol density gradient (GE Healthcare), followed by a centrifugation step for 20 minutes at 850 g at 4°C. The collected cells were washed twice in PBS (450g for 5mins, 4°C). The cells were resuspended in PBS and subsequently prepared for MS-analysis as described below.

### Flow Cytometry and Antibodies

The following antibodies were used for flow cytometry analysis and cell sorting: CD19-BV605 (SJ25-C1, BD), CD235a-APC (GA-R2, BD), CD16b-PE (CLB-gran11.5, BD), CD34-PE (581, Biolegend), CD38-APC (HIT2, Biolegend), CD45-FITC (H130, Biolegend), CD4-APC (A161A1, BioLegend), CD8-Pacific Blue (SK1, BioLegend), CD42b-FITC (HIP1, Biolegend), CD3-PE Cy7(UCHT1, Biolegend) and CD14-FITC (63D3, Biolegend). 7-Amino-Actinomycin-D (7AAD) was used as a marker for viable cells. Cells were sorted on FACS Aria II/Aria III. Collected data were analyzed using FlowJo software (Tree Star).

### Plasma preparation

Venous blood samples were centrifuged at 2,000 x g for 10 minutes at room temperature and separate aliquots of the plasma supernatants were stored at – 80°C until analysis.

### Time-resolved healthy plasma cohorts

Venous blood samples were collected from ten healthy donors at different time points as previously described[12].Informed consent was obtained in accordance with ethical permission Dnr 2013/314 LU.

### Pancreatitis Cohort

Venous blood samples from ten adult patients admitted to Skånes University Hospital and diagnosed with acute pancreatitis were included in the sample cohort. Informed consent was obtained from all patients in accordance with approved ethical permission 2009/413. Acute pancreatitis was defined as upper abdominal pain, elevated amylase levels minimum three times the upper reference limit and/or radiological findings that confirmed acute pancreatitis. Samples and centrifuged as described above. As a control, venous blood samples from 14 patients admitted to the ED at Skåne University Hospital with upper abdominal pain but with normal amylase levels were collected. Amylase analysis was performed by the Clinical Chemistry Departments at Skåne University Hospital.

### Myocardial injury cohort

Plasma samples were collected from 50 patients with chest pain admitted to the ED at Skånes University hospital, with or without increased plasma levels of troponin t. The sample collection was done unidentified at the department of clinical chemistry. Myocardial injury was considered by an elevation of cardiac troponin T values above the 99th percentile upper reference limit.

Troponin T analysis was performed by the Clinical Chemistry Departments at Skånes University Hospital Malmö on lithium heparin plasma using the Elecsys troponin T hs assay (Roche Diagnostics, Germany) run on a Cobas 6000 analyzer (Roche Diagnostics, Germany). The coefficient of variation was 2.0% both at a mean concentration of 16 ng/L and 240 ng/L.

### Infection cohort

Venous blood samples were collected from patient with suspected infection admitted to the ED at Skåne University Hospital as part of a prospective, multicenter clinical trial previously described [21]. Informed consent was obtained from all patients under an approved ethical permission (2014/741 2014-11-18). A subset of patients (n=87) with confirmed but different underlying infection (n= 40 bacterial infection + 47 viral infection) were selected for further MS-analysis. Heparin-binding protein (HBP) was analyzed in duplicate using the Axis-Shield HBP microtiter plate enzyme-linked immunosorbent assay (Axis-Shield Diagnostics). White blood cell count (WBC), C-reactive protein (CRP), and lactate analysis were performed at the Clinical Chemistry Departments at Skåne University Hospital.

### Homogenization step

The tissue biopsies were dissected into smaller pieces. Small tissue biopsies or isolated cells were mixed with 250 µl PBS (Gibco) and 100 mg 0.1 mm silica beads (Biospec Products) and homogenized using a FastPrep-96 instrument (1,600g, 180 s) (MP BIOMEDICALS). The samples were centrifuged (14,000g, 1 min, 20°C) and supernatant (referred to as SOL) was removed and stored. The remaining pellet was dissolved in 175 µl PBS + 2 % SDS (w/v), heated to 99 ° C for 5 minutes. The silica beads were removed by a new centrifugation step (14,000g, 1 min, 20°C) and the second supernatant (referred to as SDS) were removed and stored. The protein concentrations in both supernatants (SOL and SDS) were determined with Pierce BCA Protein Assay Kit (Thermo Scientific).

### SDS-PAGE analysis

Proteins from homogenized organs or cells (100 mg) were mixed with Laemmli sample buffer (Bio-Rad Laboratories Inc.) with 5% b-mercapto-ethanol (Sigma-Aldrich) and run on precasted gels (Criterion 12+2 well comb, 45 µl, Bio-Rad Laboratories Inc). The gel was run at (Criterion, Bio-Rad Laboratories Inc.) 60V until the samples started to migrate and then the voltage was increased to 160 V. The gel was subsequently stained with GelCode Blue Stain Reagent (Thermo Scientific) for 30–60 min and excessive blue stain reagent was removed using deionized water.

### In gel digestion

Proteins from homogenized organs/cells were run on SDS–PAGE followed by in gel digestion. Each lane was cut into 10 slices containing 100mM ammonium bicarbonate (ABC, Sigma-Aldrich). To destain the Gel Code Blue Stain Reagent from the gel pieces they were incubated with 50% acetonitrile (ACN, Sigma-Aldrich) 50mM ABC to shrink and then reswelled with 100mM ABC, this was repeated until no blue colour from the gel pieces could be detected. In the final washing step, the gel pieces were dehydrated using 100% ACN. The liquid was removed, and the gel pieces dried in a speedvac (miVac Duo concentrator, Genevac Ltd) before reduction with 100 mL 20mM DL-Dithiothreitol (DTT) (Sigma-Aldrich) in 100mM ABC for 60 min at 55 °C and alkylation with 200 mL 55mM iodacetamide (IAA, Sigma-Aldrich) in 100mM ABC for 45 min in the dark at room temperature. The gel pieces were washed in 100mM ABC followed by incubation in ACN. The gel pieces were then reswelled again by adding 100mM ABC and finally dehydrated by adding ACN. The liquid phase was removed, and the gel pieces completely dried in a speedvac. Proteins were digested by incubation with 100 mL trypsin (sequence grade modified trypsin porcine, 10 ng) in 100mM ABC overnight at 37 °C. Peptide were extracted by three changes of 5% formic acid in 50% ACN and by a final change of 100% ACN. The collected samples were concentrated in vacuum centrifuge and resuspended in 20 mL HPLC-water with 2% ACN, 0.2% formic acid. To all the samples, synthetic peptides (JPT Peptide Technologies, Berlin, Germany) were added for retention time calibration at the following concentrations: 1.0pM GTFIIDPGGVIR; 5.5pM TPVITGAPYEYR; 5.2pM ADVTPADFSEWSK; 2.7pM DGLDAASYYAPVR; 1.0pM GAGSSEPVTGLDAK; 2.8pM TPVISGGPYEYR; 2.9pM GTFIIDPAAVIR; 3.4pM YILAGVENSK; 2.7pM VEATFGVDESNAK; 7.7pM LGGNEQVTR. After in-gel digestion, peptides were analysed by shotgun LC-MS/MS analysis.

### SMART digest

Plasma from the different cohorts was digested with the SMART digest kit (Thermo Scientific). 1µL plasma was diluted 1:50 (v/v) with water and then further diluted 1:4 (v/v) with the SMART Digest buffer (50 mM Tris pH 7.2). Samples were transferred to SMART Digest Trypsin digest resin in PCR strips and incubated 4 h at 70°C and 1400 rpm mixing (Thermomixer C with ThermoTop, Eppendorf). Cysteines were 1) reduced by adding 2 µL 0.5M TCEP Tris(2-carboxyethyl)phosphine, (Sigma-Aldrich) and incubating 60 min at 37°C and 2) alkylated by adding 4µl 0.5M iodacetamide (Sigma-Aldrich) and incubated 60 min at room temperature. The resulting peptides were desalted using Solaµ SPE HRP plates (Thermo Scientific).

### SP3 beads

Protein solutions (supernatant from homogenized tissues and cells) were dissolved in 100 mmol/L ABC with 8 mol/L urea (Sigma-Aldrich). The proteins were reduced with TCEP (Sigma-Aldrich), final concentration 5 mM for 30 min at 37 °C and alkylated with 10 mM IAA (Sigma-Aldrich), for 30 minutes in the dark at room temperature. 10 µL of protein solution (1 µg/µl) were mixed with 2 µL beads. The bead-protein (Thermo Scientific Sera-Mag Speed Beads A and B, CAT# 4515-2105-050250, 4515-2105-050350) mixture was acidified (pH ∼2) by adding 5 µL 1% formic acid followed by adding15 µL acetonitrile (100 %) to a final concentration of 50% (v/v) of the total volume. The samples were incubated for 8 at room temperature and then placed on the magnetic stand for another 2 minutes incubation step. The supernatant was removed, followed by two rinsing steps (200 µL of 70% ethanol for 30 seconds) while the samples kept on the magnetic stand. The ethanol was removed and 180 µL of 100% acetonitrile added followed by incubation step for 15 seconds. The supernatant was discarded and the beads air dried for 30 seconds. Proteins were digested with 5 µL of 100 mM Ambic pH 8.8 that contained ∼250 ng/µL of Trypsin/rLysC enzyme mix (Promega) (Total amount 1.25 µg) over night for 14 hours at 37°C. To each sample 100% ACN was added to a final concentration of >95% After an incubation step for 8 minutes at room temperature, followed by 2 minutes on magnetic stand, the samples were washed with 180 µL 100% acetonitrile. Peptides were eluted by incubation for 5 minutes in with 20 µL 2 % DMSO, 0.1 % FA, followed by sonication and finally the samples were transferred to MS-vials along with iRT-peptides (Biognosys AG)

### LC-MS/MS analysis

**Data-dependent acquisition (DDA)** measurements for spectral library generation were performed on a Q Exactive Plus (Thermo Scientific) connected to an EASY-nLC 1000 liquid chromatography system (Thermo Scientific). Peptides were separated by C18 reverse-phase chromatography using EASY-Spray column, 25 cm x 75 µm ID, PepMap RSLC C18 2 µm with a linear gradient from 5% to 35% acetonitrile in aqueous 0.1% formic acid at a flow rate of 300 nL/min for 120 min. The 15 most intense precursor ions of charges ≥2 from an MS1 scan (R = 70,000, scan range of 400–1200 m/z) were allowed to be fragmented and measured at R = 17,500. AGC was set to 1e6 for both MS and MS/MS with ion accumulation times of 100 ms (MS) and 60 ms (MS/MS). Precursor ions were fragmented using higher-energy dissociation (HCD) at a normalized collision energy of 30.

**Data in-dependent acquisition (DIA)** measurements were performed on a Q Exactive HF-X (Thermo Scientific) connected to an EASY-nLC 1200 liquid chromatography system (Thermo Scientific). Peptides were separated by C18 reverse-phase chromatography using EASY-Spray column, 50 cm x 75 µm ID, PepMap RSLC C18 2 µm with a linear gradient from 5% to 35% acetonitrile in aqueous 0.1% formic acid at a flow rate of 300 nl/min for 120 min. The variable window data in-dependent acquisition (DIA) method is described by Bruderer et al[22]. In summary, one MS1 scan with a scan range of 350–1650 m/z, resolution of 120,000, AGC target of 3e6 and maximum IT 60 ms was followed by 26 MS2 scans with a resolution of 30,000, AGC target 3e6, auto IT and with a stepped normalized collision energy (NCE) of 25.5, 27 and 30.

MS raw data were stored and managed by openBIS (20.10.0)[23] and converted to centroided indexed mzML files with ThermoRawFileParser (1.3.1)[24].

### DIA assay library

A spectral library for all samples was built by searching both the DDA and DIA data with MSFragger-DIA using FragPipe (v18.0) following the protocol in Yu et al[25]. Peptide spectrum matches (PSMs) were validated using Percolator[26] and filtered for a 1% false discovery rate (FDR). Proteins were filtered at a 5% FDR and peptides at 1% using Philosopher (v4.4.0)[27]. The python package easypqp was then used through FragPipe with default parameters to convert the verified identifications into a spectral library for use with DIA-NN[28].

### Quantitative analysis of tissue, cell and plasma samples

DIA-NN (v1.8.1) [28]was used to analyze both the tissue and plasma DIA samples. Identifications were filtered at a 1% FDR, no protein inference was performed, smart profiling was enabled, and reanalysis of all samples using a library created from high-confidence identifications after a first-pass was enabled. The raw output from DIA-NN was passed to DPKS[29] for downstream analysis. Using DPKS, normalization was performed, and proteins were quantified using an implementation of the iq algorithm[30].

### Data Availability

Data produced in the present study are available upon reasonable request to the authors

### Assigning proteins to tissues for each atlas

Each of the six atlases was reduced to two dimensions using UMAP[31]. We then applied a Gaussian KDE for each protein, using each tissue’s relative fraction of the total ion current. Each protein-tissue pair was then projected onto a 400×400 pixel image, representing the tissues as a channel, summing the contribution for each neighbor protein. We classified each pixel by removing low-abundant channels and assigning it to a single label if one of the remaining tissues contributed more than 50% of the total signal. We would assign the pixel as common if more than three tissues contributed; otherwise, we labelled the pixel with multiple tissue labels. Last, we identified the closest pixels to each protein and used the labels for the pixel to label the protein.

### Construction of a Global distribution Atlas

The global distribution atlas combines data from all six atlases and assigns each protein to one or more tissues or cells (denoted as single or multilabel). In addition, there are two special labels: one common for proteins present in high amounts in most tissues and cells and one for primary plasma proteins.

Every protein/transcript received a weighted score from each atlas based on the underlying label assignment (UMAP/KDE classification strategy described above) and how often each analyte was identified in all atlases. We summarized the information for each protein/RNA in each tissue and cell type, weighting information from multilabel proteins proportionally to generate a global label score (GLS) ranging from 0-4. We then classified a protein as cellular, or tissue based on the amount of evidence we had for either. Once this was done, we picked the tissue or cell with the most information, including additional tissue or cell annotations, if the amount of evidence was within a pre-defined cutoff. If a protein was assigned to two or three tissue labels, they were labeled as multilabel proteins, and proteins assigned to four or more tissues were labeled as common.

To label primary plasma proteins we first selected proteins with liver labels in the RNA atlases and excluded those with an assigned liver label in the protein atlases. For this subgroup of proteins, we defined the relative abundance ratio in the liver by dividing the normalized intensities for these proteins/transcripts in the liver with the average normalized intensities across the other tissues for each tissue atlas. Comparing the results between RNA and protein atlases allowed us to identify proteins with at least a 5-fold higher abundance liver ratio in the RNA-based atlases. In the final step, we compared the selected proteins with healthy plasma proteome, identifying 114 primary plasma proteins with at least to peptides.

## Acknowledgment

E.M. is/was funded by the Wenner-Gren Foundation (FT2020-0003), Birgit and Hellmuth Hertz Foundation, Crafoord Foundation and the Swedish Society of Medicine (SLS-985287).

J.M. is a Wallenberg academy fellow (KAW 2017.0271) and is also funded by the Swedish Research Council (Vetenskapsrådet, VR) (2019-01646 and 2018-05795), the Wallenberg foundation (KAW2016.0023, KAW2019.0353 and KAW2020.0299), and Alfred Österlunds Foundation.

We want to thank Kofi Chief Nti-Karikari-Apau for the help illustrating Figur 1a.

## Notes

### Competing Interest Statement

The authors have declared no competing interest.

## References

[1] Uhlen M, Fagerberg L, Hallstrom BM, Lindskog C, Oksvold P, Mardinoglu A, et al. Tissue-based map of the human proteome. Science 2015;347:1260419–1260419. 10.1126/science.1260419.

[2] Kim M-S, Pinto SM, Getnet D, Nirujogi RS, Manda SS, Chaerkady R, et al. A draft map of the human proteome. Nature 2015;509:575–81. 10.1038/nature13302.

[3] Wilhelm M, Schlegl J, Hahne H, Gholami AM, Lieberenz M, Savitski MM, et al. Mass-spectrometry-based draft of the human proteome. Nature 2015;509:582–7. 10.1038/nature13319.

[4] Geyer PE, Holdt LM, Teupser D, Mann M. Revisiting biomarker discovery by plasma proteomics. Molecular Systems Biology 2017;13:942–15. 10.15252/msb.20156297.

[5] Thygesen K, Alpert JS, Jaffe AS, Simoons ML, Chaitman BR, White HD, et al. Third Universal Definition of Myocardial Infarction. J Am Coll Cardiol 2012;60:1581–98. 10.1016/j.jacc.2012.08.001.

[6] Meher S, Mishra TS, Sasmal PK, Rath S, Sharma R, Rout B, et al. Role of Biomarkers in Diagnosis and Prognostic Evaluation of Acute Pancreatitis. Journal of Biomarkers 2015:1–13. 10.1155/2015/519534.

[7] Malmström E, Kilsgård O, Hauri S, Smeds E, Herwald H, Malmström L, et al. Large-scale inference of protein tissue origin in gram-positive sepsis plasma using quantitative targeted proteomics. Nat Commun 2016;7:10261. 10.1038/ncomms10261.

[8] Jiang L, Wang M, Lin S, Jian R, Li X, Chan J, et al. A Quantitative Proteome Map of the Human Body. Cell 2020;183:269–283.e19. 10.1016/j.cell.2020.08.036.

[9] Moreno P, Fexova S, George N, Manning JR, Miao Z, Mohammed S, et al. Expression Atlas update: gene and protein expression in multiple species. Nucleic Acids Res 2021;50:gkab1030-. 10.1093/nar/gkab1030.

[10] Giansanti P, Samaras P, Bian Y, Meng C, Coluccio A, Frejno M, et al. Mass spectrometry-based draft of the mouse proteome. Nat Methods 2022;19:803–11. 10.1038/s41592-022-01526-y.

[11] Zhou Y, Zhou B, Pache L, Chang M, Khodabakhshi AH, Tanaseichuk O, et al. Metascape provides a biologist-oriented resource for the analysis of systems-level datasets. Nat Commun 2019;10:1523. 10.1038/s41467-019-09234-6.

[12] Stenemo M, Teleman J, Sjöström M, Grubb G, Malmström E, Malmström J, et al. Cancer associated proteins in blood plasma: Determining normal variation. PROTEOMICS 2016;16:1928–37. 10.1002/pmic.201500204.

[13] Ye X, He Y, Wang S, Wong GT, Irwin MG, Xia Z. Heart-type fatty acid binding protein (H-FABP) as a biomarker for acute myocardial injury and long-term post-ischemic prognosis. Acta Pharmacol Sin 2018;39:1155–63. 10.1038/aps.2018.37.

[14] Cao C, Li L, Zhang Q, Li H, Wang Z, Wang A, et al. Nkx2.5: a crucial regulator of cardiac development, regeneration and diseases. Front Cardiovasc Med 2023;10:1270951. 10.3389/fcvm.2023.1270951.

[15] Malmström E, Davidova A, Morgelin M, Linder A, Larsen M, Qvortrup K, et al. Targeted mass spectrometry analysis of neutrophil-derived proteins released during sepsis progression. Thrombosis and Haemostasis 2014;112. 10.1160/th14-04-0312.

[16] Bentzer P, Fisher J, Kong HJ, Mörgelin M, Boyd JH, Walley KR, et al. Heparin-binding protein is important for vascular leak in sepsis. Intensiv Care Med Exp 2016;4:33. 10.1186/s40635-016-0104-3.

[17] Singer M, Deutschman CS, Seymour CW, Shankar-Hari M, Annane D, Bauer M, et al. The Third International Consensus Definitions for Sepsis and Septic Shock (Sepsis-3). JAMA: The Journal of the American Medical Association 2016;315:801–10. 10.1001/jama.2016.0287.

[18] Habimana R, Choi I, Cho HJ, Kim D, Lee K, Jeong I. Sepsis-induced cardiac dysfunction: a review of pathophysiology. Acute Crit Care 2020;35:57–66. 10.4266/acc.2020.00248.

[19] Scott AM, Mellhammar L, Malmström E, Gustafsson AG, Bakochi A, Isaksson M, et al. Population scale proteomics enables adaptive digital twin modelling in sepsis. MedRxiv 2024:2024.03.20.24304575. 10.1101/2024.03.20.24304575.

[20] Åhrman E, Hallgren O, Malmström L, Hedström U, Malmström A, Bjermer L, et al. Quantitative proteomic characterization of the lung extracellular matrix in chronic obstructive pulmonary disease and idiopathic pulmonary fibrosis. Journal of Proteomics 2018;189:23–33. 10.1016/j.jprot.2018.02.027.

[21] Kahn F, Tverring J, Mellhammar L, Wetterberg N, Bläckberg A, Studahl E, et al. Heparin-Binding Protein as a Prognostic Biomarker of Sepsis and Disease Severity at the Emergency Department. Shock 2019;52:e135–45. 10.1097/shk.0000000000001332.

[22] Bruderer R, Bernhardt OM, Gandhi T, Xuan Y, Sondermann J, Schmidt M, et al. Optimization of Experimental Parameters in Data-Independent Mass Spectrometry Significantly Increases Depth and Reproducibility of Results*. Mol Cell Proteom 2017;16:2296–309. 10.1074/mcp.ra117.000314.

[23] Bauch A, Adamczyk I, Buczek P, Elmer F-J, Enimanev K, Glyzewski P, et al. openBIS: a flexible framework for managing and analyzing complex data in biology research. Bmc Bioinformatics 2011;12:468. 10.1186/1471-2105-12-468.

[24] Hulstaert N, Shofstahl J, Sachsenberg T, Walzer M, Barsnes H, Martens L, et al. ThermoRawFileParser: Modular, Scalable, and Cross-Platform RAW File Conversion. J Proteome Res 2020;19:537–42. 10.1021/acs.jproteome.9b00328.

[25] Yu F, Teo GC, Kong AT, Fröhlich K, Li GX, Demichev V, et al. Analysis of DIA proteomics data using MSFragger-DIA and FragPipe computational platform. Nat Commun 2023;14:4154. 10.1038/s41467-023-39869-5.

[26] The M, MacCoss MJ, Noble WS, Käll L. Fast and Accurate Protein False Discovery Rates on Large-Scale Proteomics Data Sets with Percolator 3.0. J Am Soc Mass Spectrom 2016;27:1719–27. 10.1007/s13361-016-1460-7.

[27] Leprevost F da V, Haynes SE, Avtonomov DM, Chang H-Y, Shanmugam AK, Mellacheruvu D, et al. Philosopher: a versatile toolkit for shotgun proteomics data analysis. Nat Methods 2020;17:869–70. 10.1038/s41592-020-0912-y.

[28] Demichev V, Messner CB, Vernardis SI, Lilley KS, Ralser M. DIA-NN: neural networks and interference correction enable deep proteome coverage in high throughput. Nat Methods 2020;17:41–4. 10.1038/s41592-019-0638-x.

[29] Scott AM, Hartman E, Malmström J, Malmström L. Explainable machine learning for the identification of proteome states via the data processing kitchen sink. BioRxiv 2023:2023.08.30.555506. 10.1101/2023.08.30.555506.

[30] Pham TV, Henneman AA, Jimenez CR. iq: an R package to estimate relative protein abundances from ion quantification in DIA-MS-based proteomics. Bioinformatics 2020;36:2611–3. 10.1093/bioinformatics/btz961.

[31] McInnes L, Healy J, Saul N, Großberger L. UMAP: Uniform Manifold Approximation and Projection. J Open Source Softw 2018;3:861. 10.21105/joss.00861.

