## Supplementary Figures for "Inferring the composition of the blood plasma proteome by a human proteome distribution atlas"

**Supplementary Figure 1 Malmstrom et al**

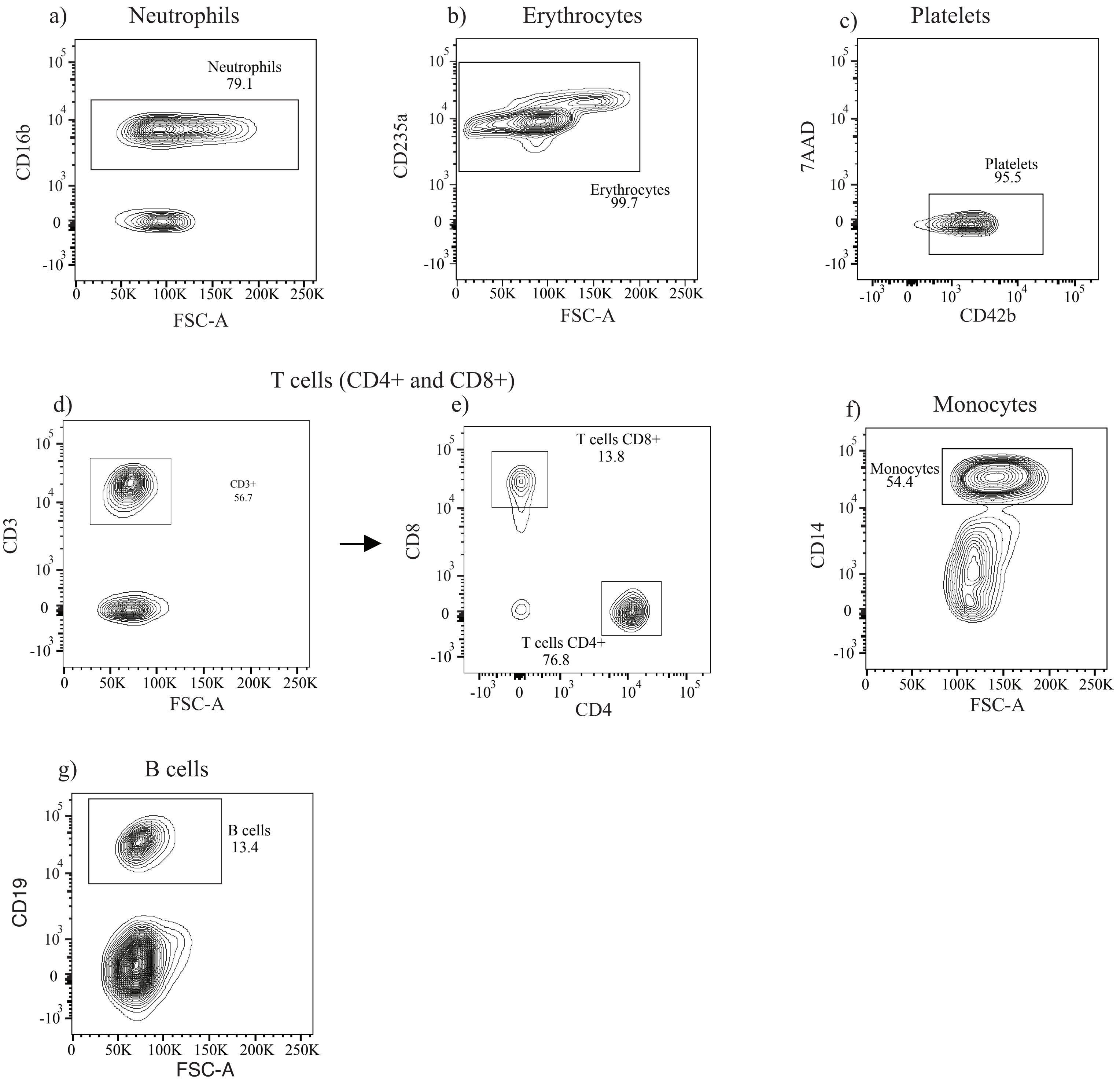

**Supplementary Figure 2 Malmstrom et al**

a)

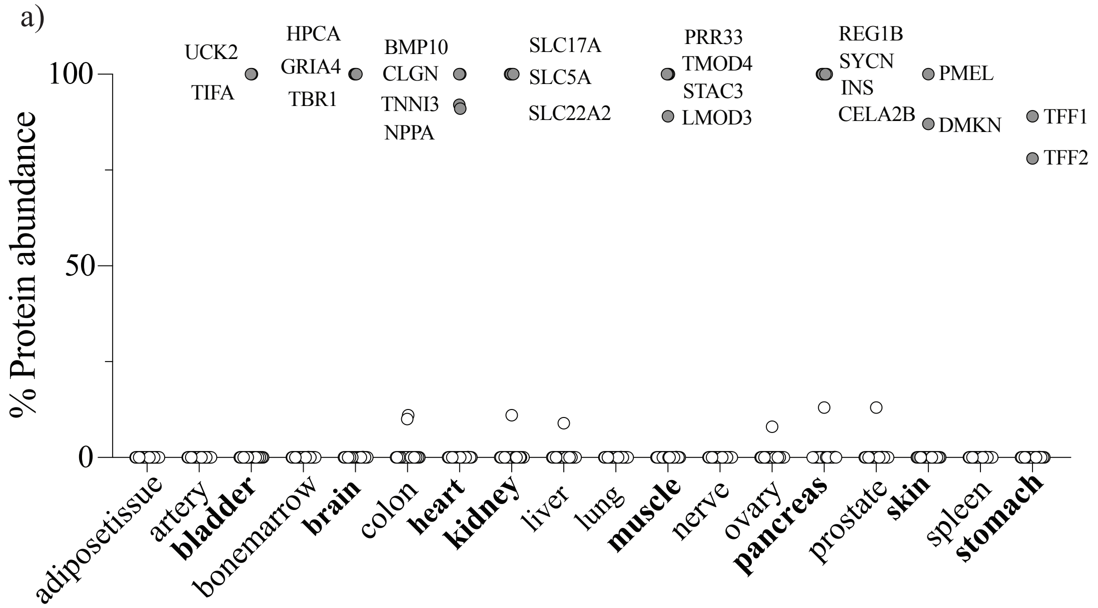


b)

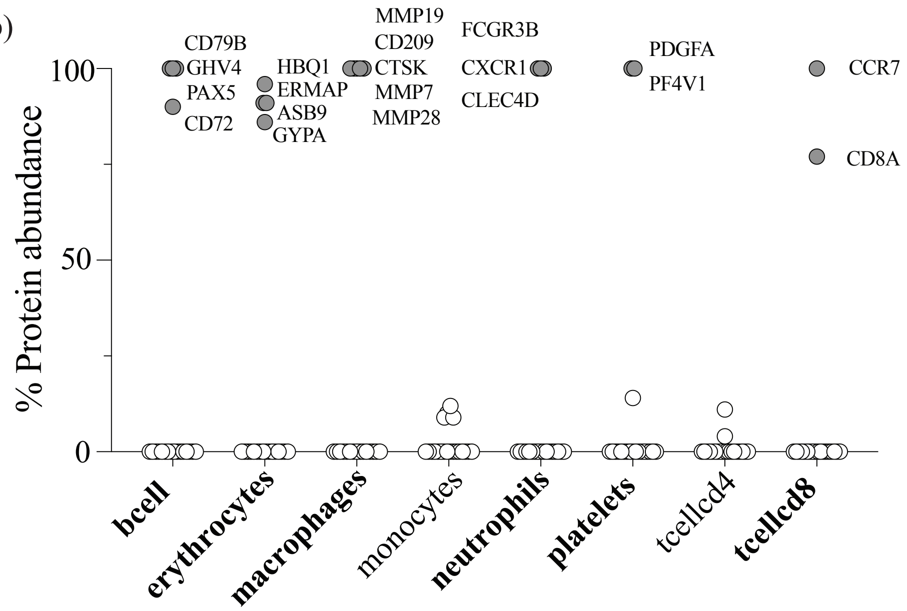

### Supplementary Figure 3 Malmstrom et al

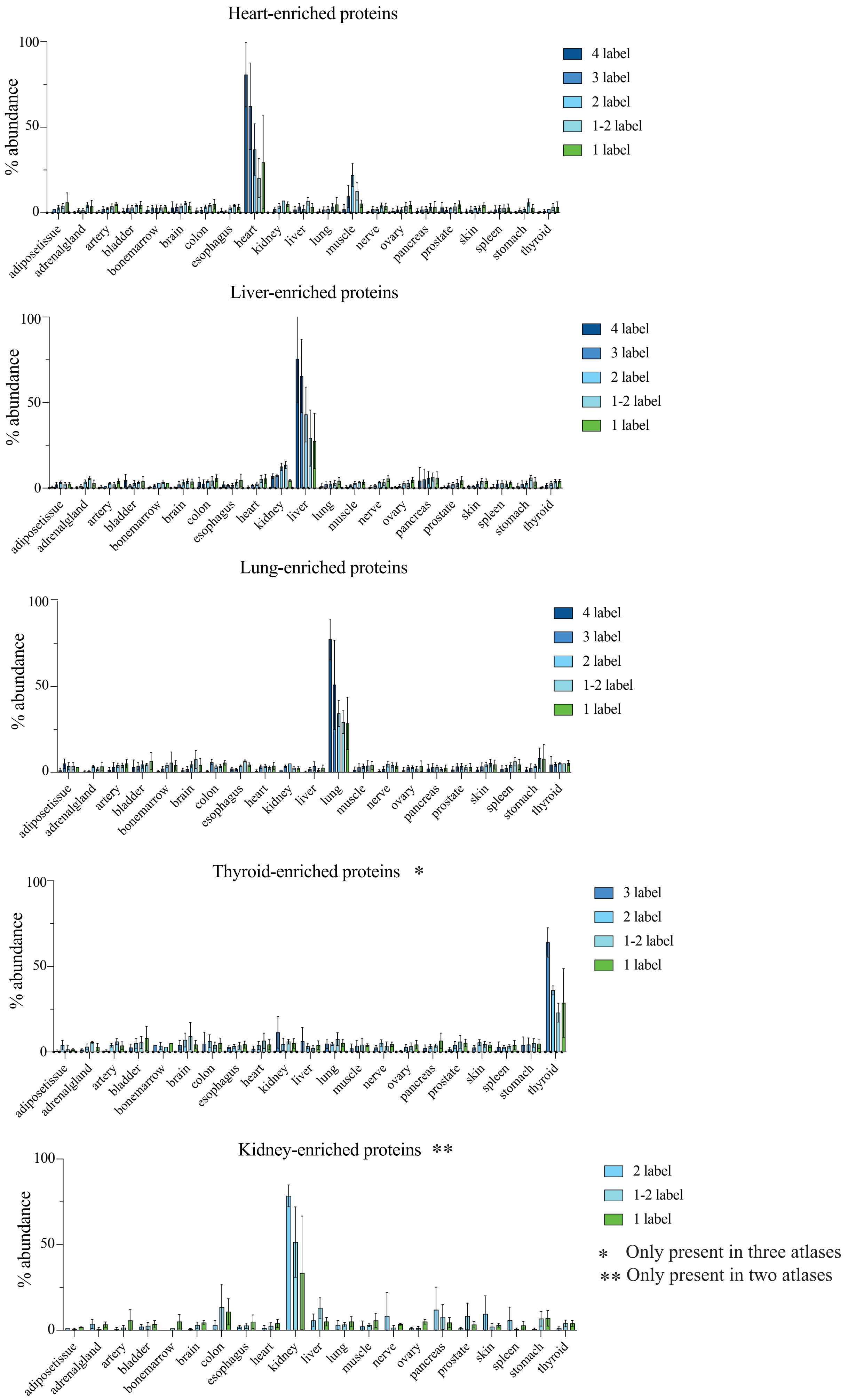

### Supplementary Figure 4 Malmstrom et al

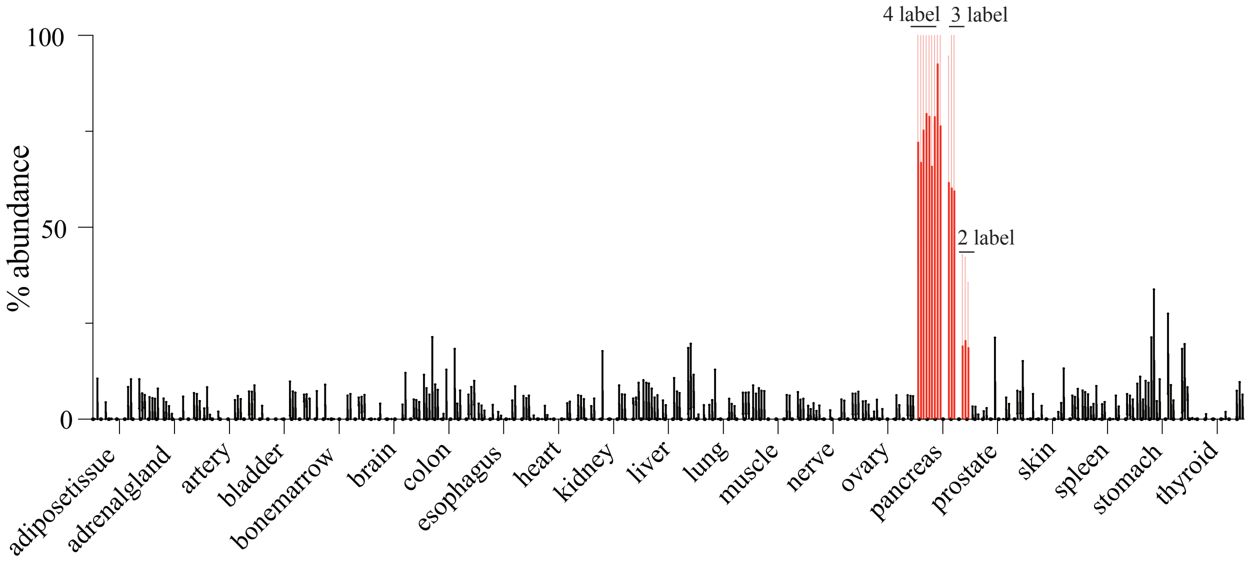

#### Supplementary Figure Legends

**Supplementary Figure 1: Outline of flow cytometry analysis and cell sorting strategy**

Images depicting the flow cytometry analysis and cell sorting strategy for the isolated blood cells.

**Supplementary Figure 2: Proteins with high tissue and cell specificity identified in HATLAS proteome atlas.**

Examples of proteins with high (A) tissue and (B) cell specificity across the HATLAS proteome atlas.

**Supplementary Figure 3: The quantitative distribution for selected tissue-enriched proteins within different levels of global label score.**

The distribution of normalized protein intensities for heart/Liver/Lung/Thyroid/Kidney-enriched proteins for each level of the global label score (GLS). The bars represent the average abundance across each atlas, the color indicates the level of GLS and error bars indicate standard deviation.

**Supplementary Figure 4: Pancreas-enriched proteins identified in the pancreatitis cohort**

Bar graph depicting the distribution of normalized protein intensities for pancreas-enriched proteins identified in pancreatitis plasma across different tissues. The bars represent the average abundance for each protein and above its GLS. Error bars indicate standard deviation.
